# Beyond paternal care: career stage and reproductive opportunities shape male services in vervet monkeys

**DOI:** 10.64898/2026.06.22.733760

**Authors:** Maria Granell-Ruiz, Josefien A. Tankink, Erica van de Waal, Carel P. van Schaik, Redouan Bshary

## Abstract

Why male primates invest in costly behaviours producing public goods remains debated, with two leading explanations, paternal care and reputation-based partner choice (RBPC). Using long-term data from four groups of wild vervet monkeys, we tested: (1) whether males show a bias in four protective “male services” (predator alarm calling, participation in between-group conflicts, leading river crossings and sentinelling); (2) which males contribute most; and (3) whether service provision predicts mating success during the mating season. We confirmed a male bias in all services. Consistent with the paternal care hypothesis, contributions were positively associated with past mating success, independently of rank, although potential fathers did not contribute more than non-fathers. Among non-fathers, service provision varied with rank, suggesting that newly immigrated males adjust their behaviour according to competitive state. Crucially, variation in alarm calling and between-group conflicts predicted future mating success, with between-group conflict emerging as the strongest and most consistent predictor of mating success across years and within mating seasons, whereas rank, tenure and social integration added little explanatory power. In contrast, sentinelling and leading river crossings did not reliably translate into mating benefits. Our findings indicate that male services are shaped by multiple selective pressures operating across different male career stages and that some forms of public goods provision function as signals of quality and cooperativeness to females. By directly linking cooperative investment to mating outcomes in a wild primate, this study provides rare empirical support for reputation-based partner choice beyond humans and highlights female choice as a potentially important force in the evolution of cooperation.

## Introduction

Cooperative acts often require individuals to invest time, energy, or risk their safety in ways that benefit others, raising the question of how such investments nevertheless enhance the donor’s fitness. Fitness returns may arise indirectly through kin selection, when helping increases the survival or reproduction of related individuals [1] or directly, when investments generate future advantages without requiring immediate reciprocity. Such direct benefits can emerge through return investments, self-serving responses of recipients, enhanced social position, or increased mating and competitive opportunities [2]. Distinguishing among these pathways is particularly challenging in the case of public goods, where a single individual’s contribution benefits multiple group members simultaneously.

One example of such public goods is the male bias in alarm calling, between-group conflict, leading group progressions and sentinelling in species where males and females form long-term associations [3–7]. These protective behaviours are cooperative actions, as they provide collective benefits while being costly or risky to the actor [3, 6–8]. Within primates, a taxon that shows above average male-female association [9], this male bias in cooperative behaviour has been found to be consistent among mating systems, a pattern referred to as “male services” [7]. This bias requires a functional explanation, as these services are costly, typically not immediately reciprocated, and occur in species in which males are the dispersing sex and are thus often unrelated to most group members [7, 10]. As a result, “male services” raise a fundamental question: what evolutionary benefits do males gain by investing in group protection despite substantial individual risk?

Several hypotheses have been proposed. Traditionally, male services have been suggested to be a form of paternal care. In this hypothesis, males that have reproduced increase their inclusive fitness by protecting the group and thereby their own offspring, in accordance with Hamilton’s rule [1, 7], which states that a helping behaviour can evolve when the cost to the actor is less than the benefits to the recipient considering their degree of relatedness. However, this hypothesis is challenged by evidence that newly immigrated males (who are unlikely to have sired offspring) also perform such services [4, 11]. Several alternative explanations have therefore been proposed. Under interdependence and social bond hypotheses, males preferentially protect particular females or their offspring, gaining future mating benefits through reciprocal relationships [12, 13]. More broadly, the group augmentation hypothesis suggests that individuals benefit directly from contributing to group defense by improving overall group survival or cohesion [14]. Another explanation is the reputation-based partner-choice (RBPC) hypothesis, which posits that males provide costly protective services to signal quality and attract mates, drawing on the handicap principle, according to which only high-quality males can afford such investments in large quantities and still survive well [8, 15, 16](for a detailed explanation of these hypotheses see [7]). Finally, male services may also function as signals in male-male competition, advertising dominance, or competitive ability to rivals rather than to females (see for an example in Arabian babblers [17]). These various hypotheses are not mutually exclusive, and different services may be shaped by different selective pressures, but simultaneous empirical testing of these hypotheses has so far been lacking.

To address this gap, we studied a population of wild vervet monkeys (*Chlorocebus pygerythrus*), which provides an ideal system for evaluating the evolutionary drivers of male services. First, they live in multimale–multifemale groups with frequent male dispersal [18–20], experience highly seasonal reproduction [21], and exhibit a mating system in which females have substantial control over their reproductive outcomes [18, 22, 23]. Second, female vervets show concealed ovulation and can actively refuse mating attempts (Dorothy L. Cheney and Seyfarth 1988; Andelman 1987) [21, 24], limiting male monopolisation and creating conditions under which male reputation and competitive signalling are likely to be important. Indeed, it appears that male rank at best modestly predicts paternity [25]. Moreover, studies from various research sites have found male vervets to overly contribute to protective behaviours compared to females, including sex-specific alarm responses to dangerous predators [26, 27], increased aggression in intergroup encounters [10, 28], a male bias in leading group progressions through risky terrain [29] and higher rates of sentinelling [3]. Besides, a previous study on male contributions to between-group conflicts (further abbreviated ‘BGC’) found support for the paternity hypothesis and some preliminary indication for the reputation-based partner choice hypothesis [10].

The behaviours considered here (alarm calling, participation in between-group conflict, leading group progressions and sentinelling) are heterogeneous in their costs and payoff structures but share a key feature: they are public behaviours that benefit multiple group members simultaneously rather than specific partners. This makes the social bonds hypothesis less likely, as support for this mechanism comes largely from partner-specific behaviours, such as male tolerance or infant protection directed at particular females. Although local proximity may influence which male initiates a given response, the resulting behaviour generally benefits the group as a whole rather than providing narrowly targeted protection. Similarly, although group augmentation may provide indirect benefits through enhanced group survival, it is less convincing in systems where females exert strong mate choice and male tenures are short. In such contexts, male cooperation is more likely to be strategic rather than an automatic by-product of group living, as males may not remain in the group long enough to benefit from delayed improvements in group viability. Thus, while other forces may also contribute, we decided to focus our analysis on disentangling the paternal care and RBPC hypotheses as the most plausible explanations in the investigated contexts. Investigating several contexts in parallel is important as different services may be shaped by different selective pressures, depending on their visibility, risk, and signalling value to females.

Assuming that we can reproduce previous studies showing the existence of male services in vervet monkeys, we furthermore tested whether male services translate into mating payoffs. We hence conducted a three-step analysis. We (1) assessed the presence of a male bias in our four behavioural contexts by testing sex differences. We (2) examined inter-male variation in the type and frequency of services provided, with the goal of understanding which males produce these services. Under the paternity hypothesis, we expected higher service investment by males with greater paternity likelihood. Under the reputation-based partner choice hypothesis, we expected service provision to covary positively with future mating opportunities, consistent with services functioning as signals to females. In that case, if services are indeed a signal of quality, we would also expect that provisioning correlates positively with male rank as an indicator of current male fighting abilities. Finally, (3) we asked what services and/or male characteristics females reward, by examining to what extent the different male services as well as other factors such as rank and tenure predict mating success during the mating season when outcomes directly impact reproductive success. By using empirical data of a multimale/multifemale primate with high levels of female choice, this study directly links male cooperative behaviour to reproductive outcomes and provides a rare empirical test of evolutionary explanations of public cooperation. In doing so, it advances our understanding of how cooperation, sexual selection and social signalling interact to shape male investment strategies in multimale societies.

## Materials and methods

### Study site and subjects

Data were collected at the INKAWU Vervet Project (IVP), located in the Mawana Game Reserve, KwaZulu-Natal, South Africa, between October 2021 and July 2025 (see table S2 for specific dates). The study focused on four habituated neighbouring groups of wild vervet monkeys. Vervet monkeys live in multimale/multifemale groups with male dispersal and female philopatry. Thus, group composition varied over time due to natural demographic processes including births, deaths, dispersals, and immigrations (Table S6).

### Data collection

Behavioural and demographic data were collected year-round by trained field assistants who passed interobserver reliability assessments and identification tests conducted regularly by the onsite scientific manager. For this study, we focused on the four core groups with the most complete datasets (AK, BD, NH and KB). We investigated four distinct types of behaviours: predator alarm calling, participation in between-group conflict, leading group progressions over rivers and time spent sentinelling.

Alarm call participation was recorded whenever an event occurred in which a predator was spotted by an observer, either terrestrial or aerial, and where there were no unidentified adult callers. In this study, we excluded encounters with snakes, as adults are less likely to approach or inspect snakes once detected [30], suggesting that they do not consider them a threat worth informing others about and thus calling would not be a potential male service. The existing primate literature supports our interpretation as there is no reported male bias in snake anti-predator behaviour [7]. We recorded participation during between-group conflict whenever two groups met each other. During these encounters, observers noted all behaviours directed towards out-group members. While some neutral or positive behaviours occurred, we focused our analyses exclusively on agonistic behaviours (Table S1). Behaviour of both groups were collected whenever trained field assistants were present in both groups. Conflicts involving unidentified adults were again excluded from the analysis. We investigated leadership during group progression. Building on a separate paper that analysed leadership in a wide range of contexts [29], we only used recordings of the first individual leading a group through potentially high-risk terrain, a river in a broad open floodplain, reaching up to more than 50 metres in width. Including river crossings (with additional data) as one of four protective services enabled direct comparison across behaviours within a unified framework. Lastly, the time spent sentinelling was recorded during focal sampling. It was defined as an individual visually scanning its surroundings from an elevated position, without visual obstruction and with an active body posture. Focal samples consisted of 20-minute continuous follows. We aimed for three focal samples on each adult individual over a 10-day period. To spread data collection and avoid biased sampling, focal observations of the same individual were never performed on the same day or in the same time block during the same 10-day period (morning, noon, afternoon).

### Variable calculation

For the first set of models, studying the potential sex-bias in cooperative behaviour, the following variables were calculated for each observation (i.e. alarm, between-group conflict, crossing and sentinelling) and individual. We calculated the proportion of males in the group, as the number of adult males divided by the total number of adults present in the group at the time of observation. To control for ecological variation, each observation was categorised into seasonal phases based on local patterns: summer (Jan–Mar), mating season (Apr–Jun), winter (Jul–Sep), and baby season (Oct–Dec). Seasons were defined based on the timing of reproductive events, with mating and birth peaks occurring approximately six months apart. These periods aligned with spring and autumn, and the remaining months were classified as summer (hot and rainy) and winter (cold and dry) based on the local temperature pattern. Due to frequent dispersal events, some males were not habituated to human observers during their first months in our study population. To address the observation bias arising from habituation differences in males, they were classified into two groups: habituated and unhabituated. Habituated males were defined as those born in an IVP-followed group or those who had spent at least 12 months in such a group.

For the second set of models, studying which males perform more cooperative behaviours, male dominance rank and social centrality were calculated from interactions recorded during the 12 months preceding each event or focal. In both cases, we used the group-standardised scores provided by each package to enable between-group comparisons. Dominance rank was derived from dyadic agonistic interactions using the EloRating package [31] (ranging from the top male, 1, to the lowest, 0), and social centrality was calculated from grooming interactions using the socialindices package [32]. Because of the promiscuity of female vervets [25] and the lack of genetic paternal data, potential paternity was estimated from tenure and its overlap with conceptions, i.e. we considered a male a potential father if he was present during the mating season when the offspring were conceived and those offspring had since been born. To assess mating success, we quantified the number of mounts recorded for each male during the year before and after each observation of a service, thereby including at least one mating season before and after the event. We did not restrict the analysis to males that were present throughout both one-year periods; thus, measurements of past and future mating success reflected the variation in male tenure within the group. Lastly, between-group conflict intensity was assigned based on the highest intensity behaviour observed during the event; if any individual exhibited high-intensity behaviour, it was classified as high, otherwise, it took the highest lower level observed (medium or low, see table S1 for classification). For group progression, we only analysed rivers where a male was found crossing first, since the male bias in leading group progression over rivers has already been established [29].

For the final model, we investigated predictors of male mating success during the mating seasons. The AK group was excluded because its territory lacks a river, preventing the calculation of leading crossings. The mating season was defined as April-June, confirmed by visual inspection of histograms of mating and births, and taking into account vervet gestation period (165 days [33]). Mating success was measured as the number of accepted mounts per male. Male services were quantified as the number of cooperative events a male participated in during the season, except for sentinelling, which was calculated as the proportion of time spent sentinelling. Since there were not enough confirmed predator encounters during the mating season, we instead recorded participation in all alarm events, including those with unknown cause, while excluding those consisting solely of snake alarm calls [27, 34]. All events with unidentified males were excluded to ensure the use of high-quality data. Rank and social integration (zCSI) were calculated on the last day of the mating season or the last day a male was present in a group. We also included tenure length (time already spent in a group) and habituation level.

### Statistical analysis

To address our three questions: (1) Are males more involved than females in protective behaviours? (2) Which males contribute the most? and (3) Which factors best predict mating success? We used two complementary modelling approaches. The first two questions were tested using a hypothesis-driven approach, fitting one model per behaviour per question using year-round data, allowing us to evaluate patterns consistent with both paternity-related and competitive hypotheses. The third question was addressed through a model-selection approach to identify which male services and social parameters best explained variation in mating success solely during the mating season, as females are receptive only at this time and direct fitness benefits through increased mating opportunities can occur only then. To simplify the manuscript, we do not present the models answering the first question (but see the model result table S3), however, they are fully documented in the GitHub repository. All analyses were conducted in R 4.4.2.

To address the second question, we investigated the predictors of male services, i.e. we asked which males contributed the most between October 2021 and July 2025. For this, we modelled participation among males as a function of Elo dominance rank, and its interactions with the possibility of paternity and the number of mounts recorded in the year before and after the observation. These models also included the proportion of males in the group, season, social centrality, habituation status, and group as fixed effects. Individual identity was included as a random intercept with rank, future and past mounts as random slopes in all models, with event identity included only when relevant. For each behaviour, we additionally fitted a null model including only control predictors and random effects and compared it to the full model using likelihood-ratio tests (table S2). All continuous predictors (e.g. Elo rank, social centrality, proportion of males, and mount counts) were standardised. For alarm calling and between-group conflict, we tested a three-way interaction between dominance rank, paternity status, and predator type or conflict intensity. Because both potential paternity and habituation level are inherently determined by tenure (see variable calculation), it was not possible to include tenure simultaneously in the models. We prioritised including these variables as they were more relevant to test our hypotheses. Lastly, to assess whether males differed consistently in their propensity to provide each service, we compared the final models used in the main analyses (Table S5), excluding random slopes, with otherwise identical models that omitted male identity as a random effect. Model comparisons were conducted using likelihood-ratio tests and changes in AIC (ΔAIC). For each service, we also extracted the variance components associated with male identity and, where relevant, event identity to quantify the magnitude of among-male and among-event variation after accounting for fixed effects.

We adopted this modelling approach for all four behaviours mentioned above. For the sentinel model, we used a negative binomial GLMM with a log-offset for observation duration and included zero-inflation terms for rank, season, and future mounts, allowing for dispersion to vary by season and the proportion of males. We excluded one outlier detected by DHARMa diagnostics, as the focal observation consisted almost entirely of sentinelling behaviour (*>* 92% of the time). All remaining behaviours were analysed as binary responses. Alarm participation was modelled with a binomial GLMM, recording whether each male called (1) or not (0) and restricting analyses to events in which at least one male vocalised. Between-group conflict participation was also modelled with a binomial GLMM, indicating whether an individual took an aggressive action toward the opposing group (1) or not (0; see list of aggressive behaviours, table S1). Group progression was analysed only in events where a male led the river crossing, recording for each male whether he was the first to cross (1) or not (0), using a negative binomial GLMM with the number of males present as an offset, since only one male per event was able to perform this behaviour. All models were fitted using the glmmTMB package [35].

Finally, to address the third question, we examined predictors of male mating success during the mating seasons of 2023 to 2025. In this way we took into account the fact that many male migration events take place just before the beginning of the mating season: any newly immigrated male cannot achieve the same total number of services than a resident male if the past year were considered, while all males are on relatively equal footing if we only consider services that happen in parallel with mating. We first fitted a global model with the number of matings (log-transformed and corrected for the number of days a male was present in that group for the season) as the response, using a linear mixed model (lmer [36]). Fixed effects included all counts of male service provision (alarm, BGC and crossing) and proportion of time spent sentinelling, dominance rank, social integration, tenure length, year, habituation status, and group identity, with male ID as a random intercept. We then performed model selection to identify which predictors best explained variation in mating success. Candidate models were compared using the dredge function in the MuMIn package [37]. Model fit was evaluated with AICc, and the relative support of each model was assessed by its ΔAICc, defined as the difference between a model and the best-fitting model. Models with ΔAICc < 2 were considered equally well supported, and summed Akaike weights (sw) were used to assess the relative importance of each predictor.

## Results

Full model specifications (Table S2) and outputs (Table S3, Table S4 and Table 1) are provided in the Supplementary Materials and GitHub repository.

**Table 1.**
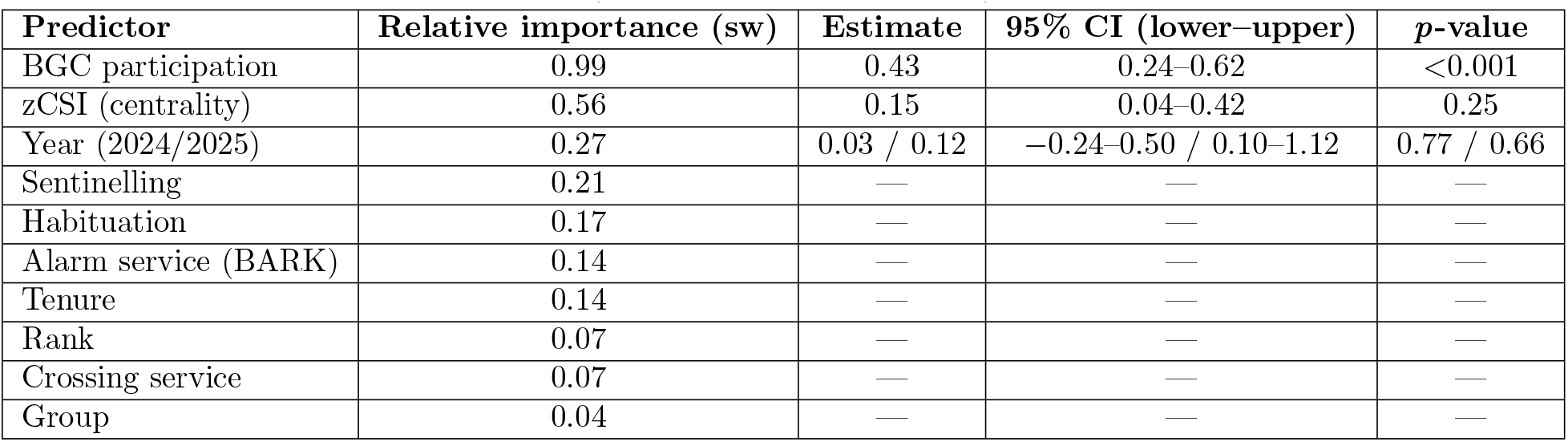
Model-averaged results for predictors of male mating success during the mating season based on the top model set (ΔAICc < 2, *n* = 3 models).

| Predictor | Relative importance (sw) | Estimate | 95% CI (lower–upper) | *p*-value |
| --- | --- | --- | --- | --- |
| BGC participation | 0.99 | 0.43 | 0.24–0.62 | <0.001 |
| zCSI (centrality) | 0.56 | 0.15 | 0.04–0.42 | 0.25 |
| Year (2024/2025) | 0.27 | 0.03 / 0.12 | –0.24–0.50 / 0.10–1.12 | 0.77 / 0.66 |
| Sentinelling | 0.21 | — | — | — |
| Habituation | 0.17 | — | — | — |
| Alarm service (BARK) | 0.14 | — | — | — |
| Tenure | 0.14 | — | — | — |
| Rank | 0.07 | — | — | — |
| Crossing service | 0.07 | — | — | — |
| Group | 0.04 | — | — | — |

We confirmed an overall male bias in all four protective behaviours that we studied, but the magnitude of the male bias varied by ecological and social contexts (Table S3, Fig. S1). Males sentinelled more (Sex:Season: *χ*^2^(3) = 16.21, *p* = 0.001, *β*_std_ = 0.166) and participated more in between-group conflict (Sex:Season: *χ*^2^(3) = 23.56, *p* < 0.001, *β*_std_ = 0.158) than females, but only during the mating season (sentinelling estimate = −0.61 ± 0.13, *p* < 0.001; BGC estimate = −0.56 ± 0.18, *p* = 0.002) and winter (sentinelling estimate = −0.55 ± 0.14, *p* < 0.001; BGC estimate = −0.79 ± 0.20, *p* < 0.001). Males also led group progressions in river crossings (*χ*^2^(1) = 12.72, *p* < 0.001, *β*_std_ = 0.472; as in Tankink et al. 2026 [29]) and called more in response to terrestrial predators (Sex:Threat: *χ*^2^(1) = 8.16, *p* = 0.004, *β*_std_ = 0.226).

Having established a male bias in these protective behaviours, we next investigated which males contributed most, to discriminate whether variation in male services can be explained by the paternal care hypothesis and/or by the RBPC hypothesis. We fitted one model per behaviour, analysing 62 predator alarm events involving 39 males, participation of 51 males in 610 between-group conflict encounters, group leading by 43 males in 75 river crossings, and sentinelling behaviour through 1,323 focal follows on 44 adult males between June 2022 and June 2024. All full models significantly differed from the null models (alarm: *χ*^2^(16) = 28.07, *p* = 0.03, ΔAIC = 3.39; between-group conflict: *χ*^2^(19) = 45.95, *p* < 0.001, ΔAIC = 8; sentinelling: *χ*^2^(11) = 35.24, *p* < 0.0001, ΔAIC = 13.3), whereas crossing differed only marginally (*χ*^2^(13) = 19.54, *p* = 0.1, ΔAIC = 6.46;Table S2).

In line with the paternal care hypothesis, both alarm calling (marginally) and between-group conflict participation were positively related to past mating success (alarm: *χ*^2^(1) = 3.69, *p* = 0.055, *β*_std_ = 0.40; between-group conflict: *χ*^2^(1) = 4.78, *p* = 0.029, *β*_std_ = 0.18; Fig. 1). There was also a weak marginal interaction between crossing events and rank (*χ*^2^(1) = 3.64, *p* = 0.06, *β*_std_ = 0.40), suggesting that males that crossed rivers more frequently tended to have fewer past matings, particularly among high-ranking males. However, the crossing result should be interpreted with caution, as none of the estimated marginal slopes were significant. There was no relationship between past mating success and time spent sentinelling (*p* = 0.40).

**Fig 1.**
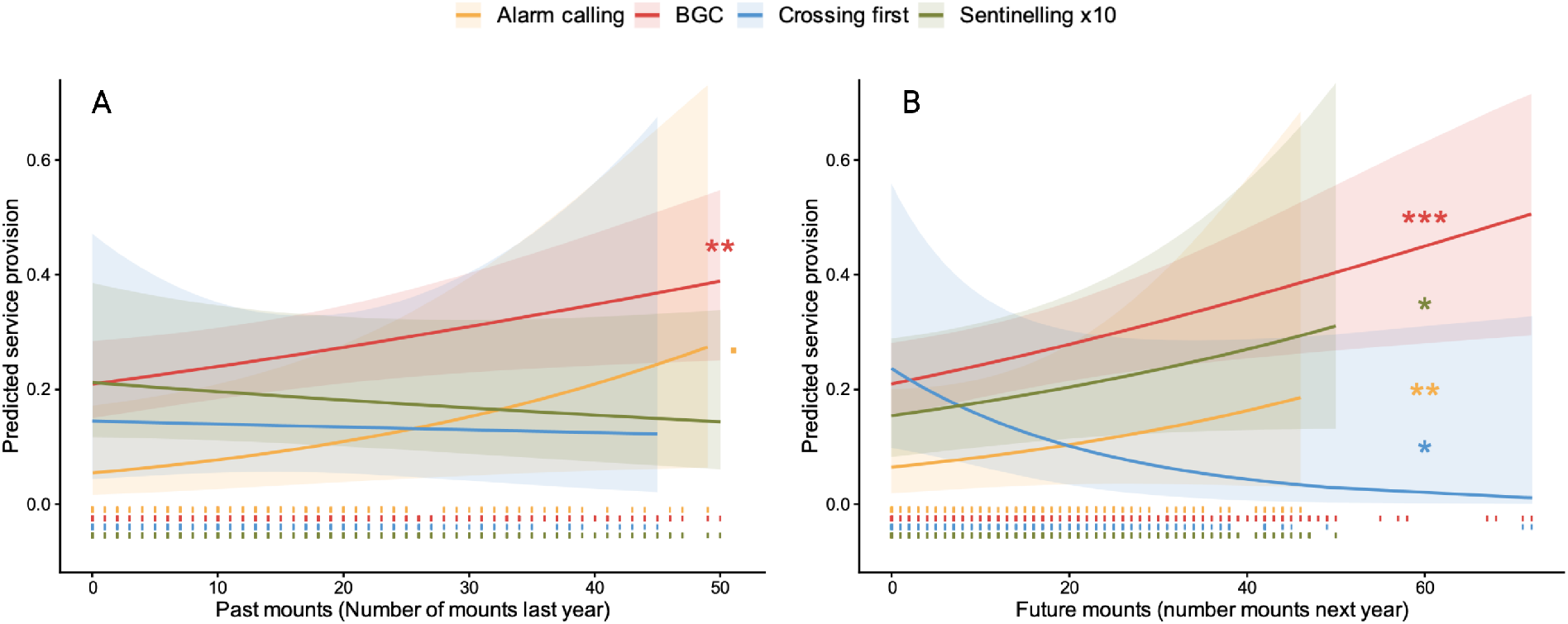
Relationship between male service provision to past (A) and future mating success (B). Each line represents a separate model prediction for a given service, and shaded bands show 95% confidence intervals. For visualization, the proportion of time spent sentinelling was multiplied by 10. Rug marks indicate the distribution of observed data points for each service. (A) Past mating success predicted participation in alarm calling and BGC only. (B) Greater participation in between-group conflict and alarm calling predicted increased mating success in the following year. Sentinelling showed a weaker positive association, whereas leading river crossings predicted reduced future mating success. Asterisks indicate statistical significance (* *p* < 0.05, ** *p* < 0.01, *** *p* < 0.001).

Contrary to the paternal care hypothesis, potential fathers did not perform more protective behaviours than non-fathers (alarm: *χ*^2^(1) = 0.10, *p* = 0.75, *β*_std_ = 0.23; sentinelling: *χ*^2^(1) = 0.167, *p* = 0.68, *β*_std_ = 0.08; crossing: *χ*^2^(1) = 0.003, *p* = 0.95, *β*_std_ = 0.46; between-group conflict: *χ*^2^(1) = 1.33, *p* = 0.72, *β*_std_ = 0.009). In general, male service rates varied with the interaction between dominance rank and potential paternity (Fig. 2). We detected significant or marginally significant effects for alarm calling (*χ*^2^(1) = 5.18, *p* = 0.023, *β*_std_ = 1.11), sentinelling (*χ*^2^(1) = 4.30, *p* = 0.038, *β*_std_ = 0.33), and leading crossings (*χ*^2^(1) = 3.82, *p* = 0.05, *β*_std_ = 0.67), but not for between-group conflict participation (*χ*^2^(1) = 0.919, *p* = 0.34, *β*_std_ = 0). When exploring conditional effects, among non-fathers, high-ranking males were more likely to alarm call (estimate = 1.10 ± 0.44, *p* < 0.001; model-predicted participation probabilities of 21% compared to 7% in low-ranking males), marginally tended to sentinel less (estimate = −0.28 ± 0.16, *p* = 0.078; 14% vs. 19%), led river crossings more often (estimate = 0.83 ± 0.38, *p* = 0.03; 3% vs. 2%), and showed a trend towards participating more in between-group conflict (estimate = −0.28 ± 0.15, *p* = 0.073; 27% vs. 24%). In contrast, among potential fathers, rank did not influence alarm calling (*p* = 0.36), leading crossings (*p* = 0.92), sentinelling (*p* = 0.145), or between-group conflict (*p* = 0.51). While participation increased with encounter intensity (*χ*^2^(2) = 111.4, *p* < 0.001), rank predicted participation only in low-intensity encounters (rank:BGC intensity: *χ*^2^(2) = 6.8, *p* = 0.03; estimate = 0.48 ± 0.15, *p* = 0.002), again driven by dominant non-fathers (estimate = 0.76 ± 0.28, *p* = 0.006), but there was no significant three-way interaction (rank:paternity status:BGC intensity: *χ*^2^(2) = 2.89, *p* = 0.23).

**Fig 2.**
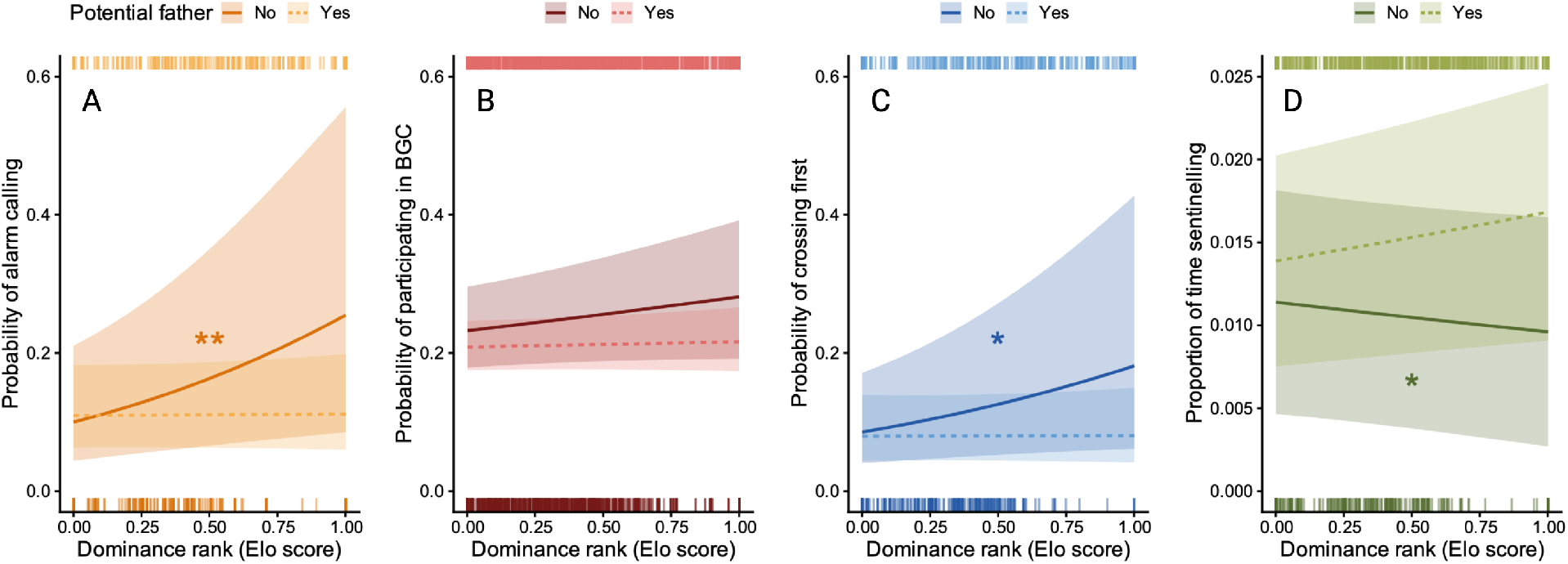
Interaction between dominance rank and paternity status on male service provision. Lines show population-level model predictions and shaded bands represent 95% confidence intervals, each panel representing one model. Top and bottom rug marks show the distribution of observed data points for potential fathers and non-fathers, respectively. Among non-fathers, higher-ranking males were more likely to alarm call (A) and initiate river crossings (C), but slightly less likely to engage in sentinelling (D). In contrast, service provision was unrelated to dominance rank in potential fathers (B). Asterisks indicate statistical significance (* *p* < 0.05, ** *p* < 0.01, *** *p* < 0.001).

Service provisioning was often associated with future mating success (Fig. 1). Males that alarm-called more during predator events and participated more in between-group conflict had more mounts in the following year (alarm: *χ*^2^(1) = 9.01, *p* = 0.003, *β*_std_ = 0.85; between-group conflict: *χ*^2^(1) = 10.88, *p* < 0.001, *β*_std_ = 0.15). This was also observed for males that sentinelled more, with a steeper increase for higher-ranking males (rank:future mounts: *χ*^2^(1) = 4.66, *p* = 0.03, *β*_std_ = 0.17; high-rank estimate = 0.30 ± 0.10, *p* = 0.001; mid-rank estimate = 0.23 ± 0.08, *p* = 0.003; low-rank estimate = 0.15 ± 0.07, *p* = 0.03) or those that sentinelled more during winter (season:future mounts: *χ*^2^(3) = 10.54, *p* = 0.015, *β*_std_ = 0.14; estimate = 0.44 ± 0.12, *p* < 0.001). Conversely, males who led river crossings surprisingly had fewer future mounts (*χ*^2^(1) = 4.36, *p* = 0.037, *β*_std_ = 0.55).

In addition to effects of dominance rank, paternity, and mating, other ecological and social factors also shaped service provision. Males sentinelled more during the mating season compared to the baby season (*χ*^2^(3) = 11.98, *p* = 0.007, *β*_std_ = 0.24; estimate = −0.48 ± 0.14, *p* = 0.002; model-predicted proportion of time spent sentinelling: 2.7% vs. 1.6%). Unhabituated individuals participated less in between-group conflict (*χ*^2^(1) = 11.68, *p* < 0.001, *β*_std_ = 0.24), while group identity explained additional variance in between-group conflict (*χ*^2^(3) = 28.91, *p* < 0.001, *β*_std_ = 0.63) and crossings (*χ*^2^(1) = 21.78, *p* < 0.001, *β*_std_ = 0.70). Despite adjusting for male numbers with an offset, the effect remained: the higher the number of males present, the less likely individuals were to lead river crossings (*χ*^2^(1) = 5.37, *p* = 0.02, *β*_std_ = 0.42), although this can be considered an obvious artefact since only one individual can be the leader. All remaining predictors were non-significant across behaviours, including the proportion of males and social integration.

We found evidence for consistent individual differences in all four services (Table S5; alarm: *χ*^2^(1) = 9.20, *p* = 0.002, ΔAIC = 7.21; BGC: *χ*^2^(1) = 62.78, *p* < 0.0001, ΔAIC = 60.8; crossing: *χ*^2^(1) = 5.05, *p* = 0.024, ΔAIC = 5.05; sentinelling: *χ*^2^(1) = 5.82, *p* = 0.016, ΔAIC = 5.09). Support for among-male differences was strongest for participation in between-group conflict, although the magnitude was only moderate (Var = 0.23, SD = 0.48) and was comparable to variation among events (Var = 0.29, SD = 0.53), showing the effect of conflict level on male participation. In contrast, alarm calling (Var = 0.66, SD = 0.81) and leading river crossings (Var = 0.54, SD = 0.73) showed the greatest among-male variation, while event-level variation was negligible. Sentinelling exhibited the weakest among-male differences (Var = 0.065, SD = 0.25). Together, these results indicate that some males consistently contributed more than others across all services.

To evaluate whether some services are stronger predictors of imminent mount success than others, we next modelled service provision specifically during the mating season. Model selection identified three models within ΔAICc < 2, all including between-group conflict participation as a key predictor. Model averaging confirmed that between-group conflict participation was the strongest predictor of mating success (estimate = 0.43, 95% CI: 0.24–0.62, *p* < 0.001; sw = 0.99;Fig. 3), with males participating more in group protection having more mounts. Males more integrated within the group also tended to have more mounts, but this effect was weaker and non-significant (sw = 0.56, *p* = 0.25). There was also non-significant variation between years (sw = 0.27, *p >* 0.66), and all remaining predictors, including rank, tenure, and other service types, had low relative importance (sw < 0.21; Table 1). Overall, participation in between-group conflict during the mating season best predicted mating success within the same period.

**Fig 3.**
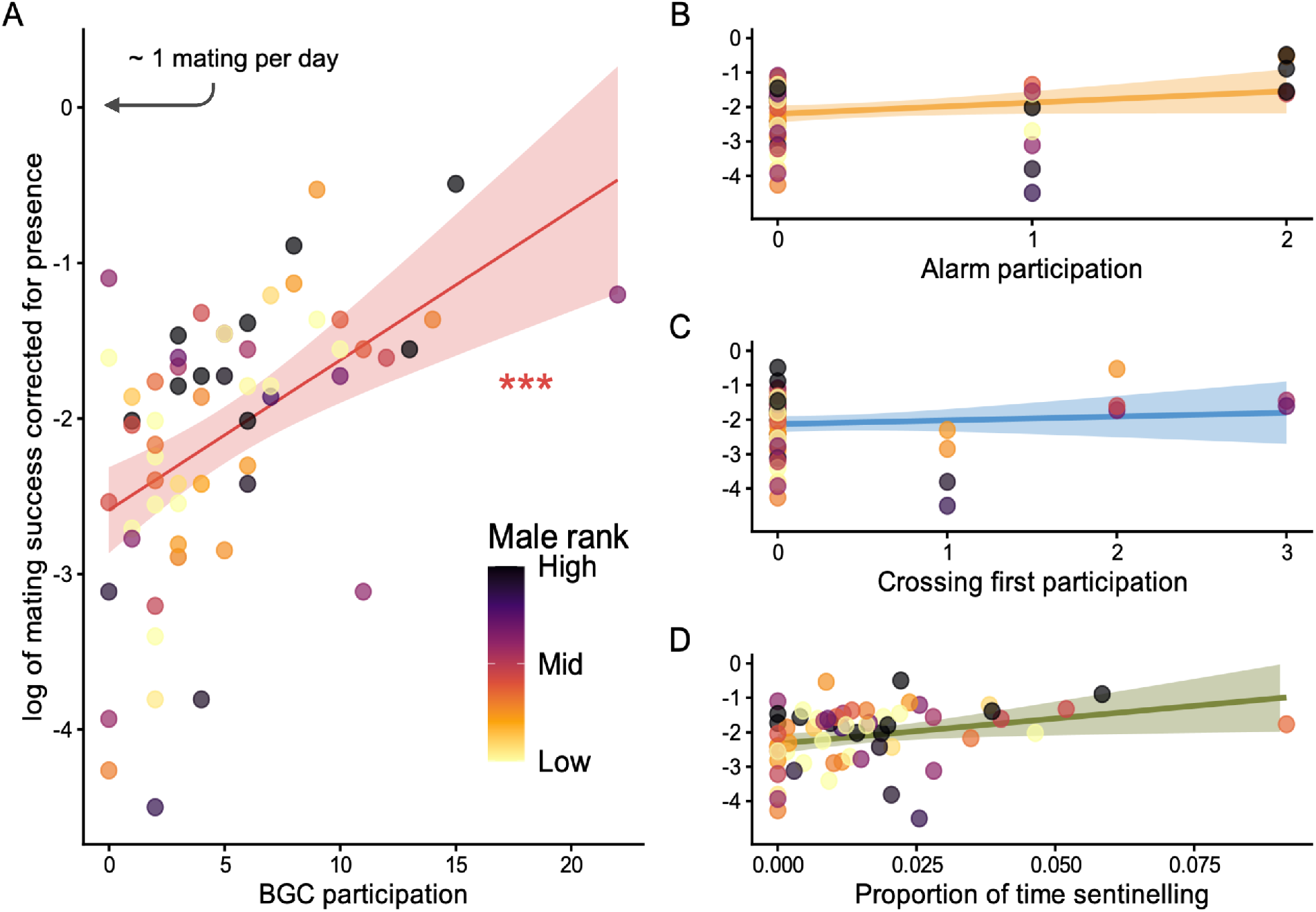
Relationship between male services and mating rate (log-transformed and corrected for days present) during the mating season. Values around 0 correspond to approximately one mating per day, indicating very high performance. Points represent raw observations for individual males and are coloured by dominance rank. Panel A shows participation in between-group conflicts; the solid line indicates model predictions with 95% confidence intervals from the best-supported model, with increased participation associated with higher mating success (*** *p* < 0.001), independent of rank. Panels B–D show the number of alarm-calling events, river crossings led, and the proportion of time spent sentinelling, respectively. Lines and confidence intervals in these panels are based on the raw data, as these variables were not retained in the best-supported model.

Relative importance (sw) reflects the summed Akaike weights across all candidate models. Estimates, 95% confidence intervals, and *p*-values are derived from model averaging across the top model set. Predictors not retained in the best-supported models are indicated with dashes. Between-group conflict participation emerged as the strongest predictor of mating success, whereas all other predictors received comparatively little support.

## Discussion

This study aimed to identify the drivers of male service provisioning in vervet monkeys, a species with a high degree of female choice. Consistent with previous work, we confirmed a male bias in service provisioning [3, 7, 10, 29], where males were more likely than females to lead river crossings and respond to terrestrial predators, as well as to sentinel and participate in between-group conflict, particularly during the mating and winter seasons. The relation between rank and service provisioning depended on male career stage, with the relationship being flat for potential fathers and positively or negatively correlated in newly immigrated males. Our study further shows that part of the variation in male service provisioning is predicted by past and future mating success. Thus, mating success emerges as a potential key currency that females offer males for their services [38]. However, these benefits are unevenly distributed across services and male career stages when entering a group. Across all behaviours examined, participation in between-group conflict emerged as the most consistent and strongest correlate of mating success, both across years and within the mating season itself. Alarm calling was also positively correlated with mating success, whereas sentinelling showed a weak and inconsistent trend, and leading river crossings did not translate into mating benefits. This pattern suggests that the four public services are not equally salient to females and hence not equally effective as quality signals. We discuss the implications of these results below.

### Male service and male career stage

Variation in male service provisioning was largely explained by an interaction between paternity status and position in the hierarchy. We refer to the combination of these two variables as defining a male’s career stage [10, 39]. New immigrants and hence non-fathers adjust their alarm, crossing and sentinelling services to their rank. Dominant non-fathers alarm called more, crossed rivers first more often and were the main participants in low-intensity between-group encounters, while spending less time sentinelling. This suggests that new immigrants play a competitive reputation game, targeting both females to increase future mating success and males as to establish and strengthen their dominance. The RBPC hypothesis predicts that performance in such competition reflects individual quality and state [13, 16], this has often been proxied by dominance rank in primates. We note, however, that rank effects on service provisioning were not particularly strong. In vervet monkeys, female influence on male hierarchies [18, 23] may weaken the link between rank and fighting ability. In this context, rank may reflect a transient competitive state rather than quality, especially in newly immigrated males, who might establish themselves through elevated service provisioning.

In contrast, potential fathers set a level of service provisioning that is rather independent of their rank (but correlated with past mating, see below). Thus, it appears that high-ranking potential fathers do not signal their supposed inherent or current quality to females or to other males. This is in line with the hypothesis that once males have secured mating opportunities, the benefits of escalating costly displays diminish, while risk and opportunity cost increase, even for dominant males [39–41]. Conversely, low-ranking potential fathers apparently do not upregulate sentinelling to monitor rivals. Thus, it appears that potential fathers stop ‘playing games’ but instead do ‘what fathers do’.

### Male services and mating success

Of the four male services, participation in between-group conflict, and to a lesser extent predator alarm calling, were positively associated with both future and past mating success. The results suggest these two behaviours are under selection through benefits that are directly linked to mating. On one hand, males appear to adjust their contributions based on past mating success, which might serve as a proxy for paternity probability. This pattern is consistent with the paternity hypothesis, a scenario in which males that contribute more are the ones that have the higher chance to have offspring. On the other hand, males may provide these public services to increase their future matings, which is in line with predictions from the reputation-based partner choice hypothesis [16]. Notably, we did not find a relationship between mating success and dominance rank. Thus, to achieve more matings, males apparently have to build their reputation by service provisioning rather than by dominance. As rank is often related to body condition and competitive ability in primates, it is rather unclear why high-ranking males do not consistently participate more in intergroup encounters, as one would predict based on the honest signalling hypothesis [15, 16]. While vervets’ hierarchies change frequently due to frequent dispersal and female intervention [18, 23], a positive correlation between rank and matings would be expected as long as females support the strongest or the best service providers. An alternative consideration of the RBPC framework, in alignment with other game theory models, is that variation in service provisioning might be driven by differences in individual willingness or ability to contribute [16]. Consistent with this prediction, male identity explained significant variation in all four services, particularly in BGC, indicating that some males are more inclined or able to provide services than others. Thus, while males can apparently increase their mating success through contributions to between-group conflicts, and to a lesser extent to non-snake predator alarm calling, the causes underlying male variation in service provisioning may reflect variation in quality, willingness to contribute, or both.

The importance of male participation in between-group conflicts for female choice suggests that females value this service more than others, consistent with the observation that female reproductive success is highly dependent on access to food and hence territory quality [10, 34]. Accordingly, previous research showed that female vervets closely monitor male behaviour during such conflicts, actively recruiting males to participate and rewarding participants with grooming [42].

### Male services not linked to mating success

Two of the four studied male services, sentinelling and leading river crossings, were not reliably positively associated with mating success. Thus, it appears unlikely that these two services are under positive selection as a form of paternal care or a way to increase male reputation in the eyes of females. Indeed, alternative explanations exist for the role of sentinelling behaviour. In vervets, sentinelling has been described as self-serving behaviour, with individuals monitoring social competitors rather than predators [3]. The observed increased sentinelling among low-ranking males may hence reflect social monitoring rather than cooperative behaviour [43, 44]. The negative results on river crossings were unexpected, given that leading progressions is indeed risky [7]. As females apparently do not care about males leading in river crossings, other selective pressures might be at play. For instance river crossings may function in male–male signalling contexts, consistent with the idea that some conspicuous and risky acts are shaped primarily by intrasexual competition rather than female preference [17, 45]. However, a recent study on the same population found no male rank increases related to leading rates [29]. This result does not necessarily exclude a signalling function, as leadership was already biased towards high-ranking males, potentially limiting opportunities for further rank increases. Although we did not test this hypothesis directly and it therefore remains speculative, future studies should investigate whether leading risky group progressions yields benefits that are not captured by mating success alone [46, 47]. Taken together, these findings suggest that some male services may be influenced by status maintenance or male–male competitive dynamics rather than female choice, although further research is needed to evaluate these possibilities.

## Conclusion

In conclusion, our results suggest that male services are partly shaped by direct offspring defence but also, and perhaps more so, by reputation-based partner choice, with both processes operating differently across male career stages. More broadly, these findings contrast with classic public-goods and cooperation models that often assume homogeneous players and uniform payoffs. Instead, the results support approaches in which cooperative investment is sex-specific, state-dependent and strategically targeted [38, 48, 49].

Importantly, cooperative acts may have multiple targets: males may direct some services primarily toward own offspring and females, yet other group members (including rivals and juveniles) benefit as a by-product, and male–male competition may drive public goods production even when females do not incorporate such contributions in their assessment of males. Empirical support for reputation-based partner choice outside humans has been limited (in humans: [50–53]), yet by linking public cooperative behaviour to mating outcomes in a non-human primate our study provides evidence that such dynamics operate beyond humans. Future work should determine whether our correlate of fitness, i.e. mating frequency, indeed translates into actual paternity. Altogether, these findings highlight the role of female choice in shaping male cooperation, suggesting that protective services can operate as strategic extensions of mating effort. Such dynamics may be widespread in social species and deserve broader attention in cooperation theory.

## Acknowledgments

We are thankful to the iNkawu Vervet Project team for their dedication to long-term group monitoring, data collection, and logistical support throughout this study. Special thanks go to Nokubonga Dhlamini, Michael Henshall, Siboniso Thela, and Zonke Mbutho for managing the field site and providing invaluable assistance in the field. We are grateful to the van der Walt family for allowing us to conduct research on their reserve and to Ezemvelo KZN Wildlife for issuing the permits required for this work. We also thank Radu Slobodeanu and Roger Mundry for their statistical advice and support. This research was funded by the Swiss National Science Foundation (grant 310030 197884). Fieldwork expenses were supported through grants awarded to EvdW by the Swiss National Science Foundation (PP00P3 198913), the ProFemmes programme of the Faculty of Biology and Medicine at the University of Lausanne, and the European Research Council through the Horizon 2020 research and innovation programme (ERC Starting Grant “KNOWLEDGE MOVES”, grant agreement No. 949379).

## References

1. Hamilton WD. The genetical evolution of social behaviour. II. J Theor Biol. 1964;7(1):17–52. doi:10.1016/0022-5193(64)90039-6.

2. Bshary R, Bergmüller R. Distinguishing four fundamental approaches to the evolution of helping. J Evol Biol. 2008 Mar;21(2):405–20. doi:10.1111/j.1420-9101.2007.01482.x.

3. Baldellou M, Peter Henzi S. Vigilance, predator detection and the presence of supernumerary males in vervet monkey troops. Animal Behaviour. 1992 Mar;43(3):451–61. doi:10.1016/s0003-3472(05)80104-6.

4. Stephan C, Zuberbühler K. Persistent Females and Compliant Males Coordinate Alarm Calling in Diana Monkeys. Current Biology. 2016 Nov;26(21):2907–12. doi:10.1016/j.cub.2016.08.033.

5. Stojan-Dolar M, Heymann EW. Vigilance in a Cooperatively Breeding Primate. Int J Primatol. 2010 Feb;31(1):95–116. doi:10.1007/s10764-009-9385-7.

6. Van Schaik CP, Van Noordwijk MA. The special role of male Cebus monkeys in predation avoidance and its effect on group composition. Behav Ecol and Sociobiology. 1989 May;24(5):265–76. doi:10.1007/bf00290902.

7. Van Schaik CP, Bshary R, Wagner G, Cunha F. Male anti-predation services in primates as costly signalling? A comparative analysis and review. Ethology. 2022 Jan;128(1):1–14. doi:10.1111/eth.13233.

8. Roberts G. Competitive altruism: from reciprocity to the handicap principle. Proceedings of the Royal Society of London Series B: Biological Sciences. 1998 Mar;265(1394):427–31. doi:10.1098/rspb.1998.0312.

9. Ostner J, Vigilant L, Bhagavatula J, Franz M, Schülke O. Stable heterosexual associations in a promiscuous primate. Anim Behav. 2013;86(3):623–31. doi:10.1016/j.anbehav.2013.07.004.

10. Arseneau TJM, Taucher AL, van Schaik CP, Willems EP. Male monkeys fight in between-group conflicts as protective parents and reluctant recruits. Animal Behaviour. 2015 Dec;110:39–50. doi:10.1016/j.anbehav.2015.09.006.

11. Matsumoto-Oda A, Okamoto K, Takahashi K, Ohira H. Group size effects on inter-blink interval as an indicator of antipredator vigilance in wild baboons. Scientific Reports. 2018;8(1):10062. doi:10.1038/s41598-018-28174-7.

12. Eshel I, Shaked A. Partnership. J Theor Biol. 2001 Feb;208(4):457–74. doi:10.1006/jtbi.2000.2232.

13. Roberts G. Cooperation through interdependence. Anim Behav. 2005;70(4):901–8. doi:10.1016/j.anbehav.2005.02.006.

14. Kokko H, Johnstone RA, Th Cb. The evolution of cooperative breeding through group augmentation. Proceedings of the Royal Society of London Series B: Biological Sciences. 2001;268(1463):187–96. doi:10.1098/rspb.2000.1349.

15. Zahavi A. Mate selection—a selection for a handicap. J Theor Biol. 1975;53(1):205–14. doi:10.1016/0022-5193(75)90111-3.

16. Roberts G, Raihani N, Bshary R, Manrique HM, Farina A, Samu F, et al. The benefits of being seen to help others: indirect reciprocity and reputation-based partner choice. Philos Trans R Soc Lond B Biol Sci. 2021 Nov;376(1838):20200290. doi:10.1098/rstb.2020.0290.

17. Dattner A, Zahavi A, Zahavi A. Competition over guarding in the Arabian babbler (Turdoides squamiceps), a cooperative breeder. F1000Research. 2016;4:618. doi:10.12688/f1000research.6739.2.

18. Young C, McFarland R, Barrett L, Henzi SP. Formidable females and the power trajectories of socially integrated male vervet monkeys. Animal Behaviour. 2017 Mar;125:61–7. doi:10.1016/j.anbehav.2017.01.006.

19. Henzi SP, Lucas JW. Observations on the Inter-Troop Movement of Adult Vervet Monkeys (Cercopithecus aethiops). Folia Primatol. 1980 Dec;33(3):220–35. doi:10.1159/000155936.

20. Isbell LA, Van Vuren D. Differential Costs of Locational and Social Dispersal and Their Consequences for Female Group-Living Primates. Behaviour. 1996;133(1-2):1–36. doi:10.1163/156853996X00017.

21. Cheney DL, Seyfarth RM. Assessment of meaning and the detection of unreliable signals by vervet monkeys. Animal Behaviour. 1988 Apr;36(2):477–86. doi:10.1016/S0003-3472(88)80018-6.

22. Keddy AC. Female mate choice in vervet monkeys (Cercopithecus aethiops sabaeus). Am J Primatol. 1986 Jan;10(2):125–34. doi:10.1002/ajp.1350100204.

23. Hemelrijk CK, Wubs M, Gort G, Botting J, Van De Waal E. Dynamics of Intersexual Dominance and Adult Sex-Ratio in Wild Vervet Monkeys. Frontiers in Psychology. 2020 May;11. doi:10.3389/fpsyg.2020.00839.

24. Andelman SJ. Evolution of concealed ovulation in vervet monkeys (Cercopithecus aethiops). The American Naturalist. 1987;129(6):785–99.

25. Minkner MM, Young C, Amici F, McFarland R, Barrett L, Grobler JP, et al. Assessment of male reproductive skew via highly polymorphic STR markers in wild vervet monkeys, Chlorocebus pygerythrus. Journal of Heredity. 2018;109(7):780–90. doi:10.1093/jhered/esy048.

26. Seyfarth RM, Cheney DL, Marler P. Monkey Responses to Three Different Alarm Calls: Evidence of Predator Classification and Semantic Communication. Science. 1980 Nov;210(4471):801–3. doi:10.1126/science.7433999.

27. Schad L, Van De Waal E, Fischer J. Anti-Snake Behavior and Snake Discrimination in Vervet Monkeys. Ethology. 2025 Mar;131(3). doi:10.1111/eth.13541.

28. Cheney DL. Intergroup Encounters among Free-Ranging Vervet Monkeys. Folia Primatol. 1981 Jan;35(2-3):124–46. doi:10.1159/000155970.

29. Tankink JA, van de Waal E, Bshary R, van Schaik CP. Leadership Under Risk: Male Vervet Monkeys’ (Chlorocebus pygerythrus) Roles in Group Progression Across Potentially High-Risk Terrain. Ethology. 2026. doi:10.1111/eth.70085.

30. Schad L, Dongre P, Van De Waal E, Fischer J. Loud Call Production in Male Vervet Monkeys (Chlorocebus pygerythrus) Varies with Season and Signaller Rank. Int J Primatol. 2025 Apr;46(2):538–55. doi:10.1007/s10764-024-00475-x.

31. Neumann C, Kulik L. EloRating: Animal dominance hierarchies by Elo rating. The R Foundation; 2014. R package version accessed June 2026. https://github.com/cran/EloRating. doi:10.1016/j.anbehav.2011.07.016.

32. Neumann C. socialindices: Calculate Sociality Indices from Animal Behaviour Data. The R Foundation; 2016. R package version accessed June 2026. https://github.com/gobbios/socialindices.

33. Rowell T. Reproductive cycles of two Cercopithecus monkeys. Reproduction. 1970;22(2):321–38.

34. Bshary R, Richter XYL, van Schaik C. Male services during between-group conflict: the ‘hired gun’ hypothesis revisited. Philos Trans R Soc Lond B Biol Sci. 2022 Apr;377(1851):20210150. doi:10.1098/rstb.2021.0150.

35. Brooks ME, Kristensen K, van Benthem KJ, Magnusson A, Berg CW, Nielsen A, et al. glmmTMB Balances Speed and Flexibility Among Packages for Zero-inflated Generalized Linear Mixed Modeling. The R Journal. 2017;9(2):378–400. doi:10.32614/RJ-2017-066.

36. Bates D, Mächler M, Bolker B, Walker S. Fitting Linear Mixed-Effects Models Using lme4. Journal of Statistical Software. 2015;67(1):1–48. doi:10.18637/jss.v067.i01.

37. Bartoń K. MuMIn: Multi-Model Inference; 2026. R package version 1.48.19. https://CRAN.R-project.org/package=MuMIn.

38. Sterck EHM, Crockford C, Fischer J, Massen JJM, Tiddi B, Perry S, et al. The evolution of between-sex bonds in primates. Evol Hum Behav. 2024 Nov;45(6): doi:10.1016/j.evolhumbehav.2024.106628.

39. van Noordwijk MA, van Schaik CP. Sexual selection and the careers of primate males: paternity concentration, dominance–acquisition tactics and transfer decision. Sexual selection in primates: New and comparative perspectives. 2004:208–29 doi:10.1006/jtbi.2000.2093.

40. Kokko H. Evolutionarily stable strategies of age-dependent sexual advertisement. Behav Ecol and Sociobiology. 1997 Aug;41(2):99–107 doi:10.1007/s002650050369.

41. Silk JB, Städele V, Roberts EK, Vigilant L, Strum SC. Shifts in Male Reproductive Tactics over the Life Course in a Polygynandrous Mammal. Current Biology. 2020 May;30(9):1716–20.e doi:10.1016/j.cub.2020.02.013.

42. Arseneau-Robar TJM, Taucher AL, Müller E, Van Schaik C, Bshary R, Willems EP. Female monkeys use both the carrot and the stick to promote male participation in intergroup fights. Proc R Soc B. 2016 Nov;283(1843): doi:10.1098/rspb.2016.1817.

43. Gosselin-ildari AD, Koenig A. The Effects of Group Size and Reproductive Status on Vigilance in Captive Callithrix jacchus. Am J Primatol. 2012 Jul;74(7):613–21 doi:10.1002/ajp.22013.

44. Allan ATL, Hill RA. What have we been looking at? A call for consistency in studies of primate vigilance. American Journal of Physical Anthropology. 2018 Feb;165(S65):4–22 doi:10.1002/ajpa.23381.

45. Gintis H, Smith EA, Bowles S. Costly Signaling and Cooperation. J Theor Biol. 2001 Nov;213(1):103–19 doi:10.1006/jtbi.2001.2406.

46. Hawkes K, Bliege Bird R. Showing off, handicap signaling, and the evolution of men’s work. Evolutionary Anthropology: Issues, News, and Reviews. 2002 Jan;11(2):58–67 doi:10.1002/evan.20005.

47. Smith KM, Apicella CL. Partner choice in human evolution: The role of cooperation, foraging ability, and culture in Hadza campmate preferences. Evol Hum Behav. 2020;41(5):354–66 doi:10.1016/j.evolhumbehav.2020.07.009.

48. Mcnamara JM, Székely T, Webb JN, Houston AI. A dynamic game-theoretic model of parental care. J Theor Biol. 2000;205(4):605–23 doi:10.1006/jtbi.2000.2093.

49. Leimar O. Life-history analysis of the Trivers and Willard sex-ratio problem. Behav Ecol. 1996;7(3):316–25 doi:10.1093/beheco/7.3.316.

50. Farrelly D, Lazarus J, Roberts G. Altruists Attract. Evolutionary Psychology. 2007 Apr;5(2): doi:10.1177/147470490700500205.

51. Iredale W, Van Vugt M, Dunbar R. Showing Off in Humans: Male Generosity as a Mating Signal. Evolutionary Psychology. 2008 Jul;6(3): doi:10.1177/147470490800600302.

52. Barclay P. Altruism as a courtship display: Some effects of third-party generosity on audience perceptions. British Journal of Psychology. 2010 Feb;101(1):123–35 doi:10.1348/000712609X435733.

53. Raihani N, Smith S. Competitive Helping in Online Giving. Current Biology. 2015 May;25(9):1183–6 doi:10.1016/j.cub.2015.02.042.

