## Supplementary materials for "Beyond paternal care: career stage and reproductive opportunities shape male services in vervet monkeys"

Table S1: BGC score system

| Category | Score | Behaviours |
| --- | --- | --- |
| high | 3 | “Attack", "Fight", "Bite", "Chase", "Face offs", "face-off" |
| medium | 2 | "Advance fast", "Front line (ind at the interface)", "Stare", "Head bob" |
| minor | 1 | "Advance slow", "Contact calls", "Alarm calls", "Vocalise", "Vigilant", "Chorus calls", "Stand-up", "aggression call" |

**Table S1**. Score system used to weight the behaviours performed during between-group conflict and determine the intensity of individuals’ participation.

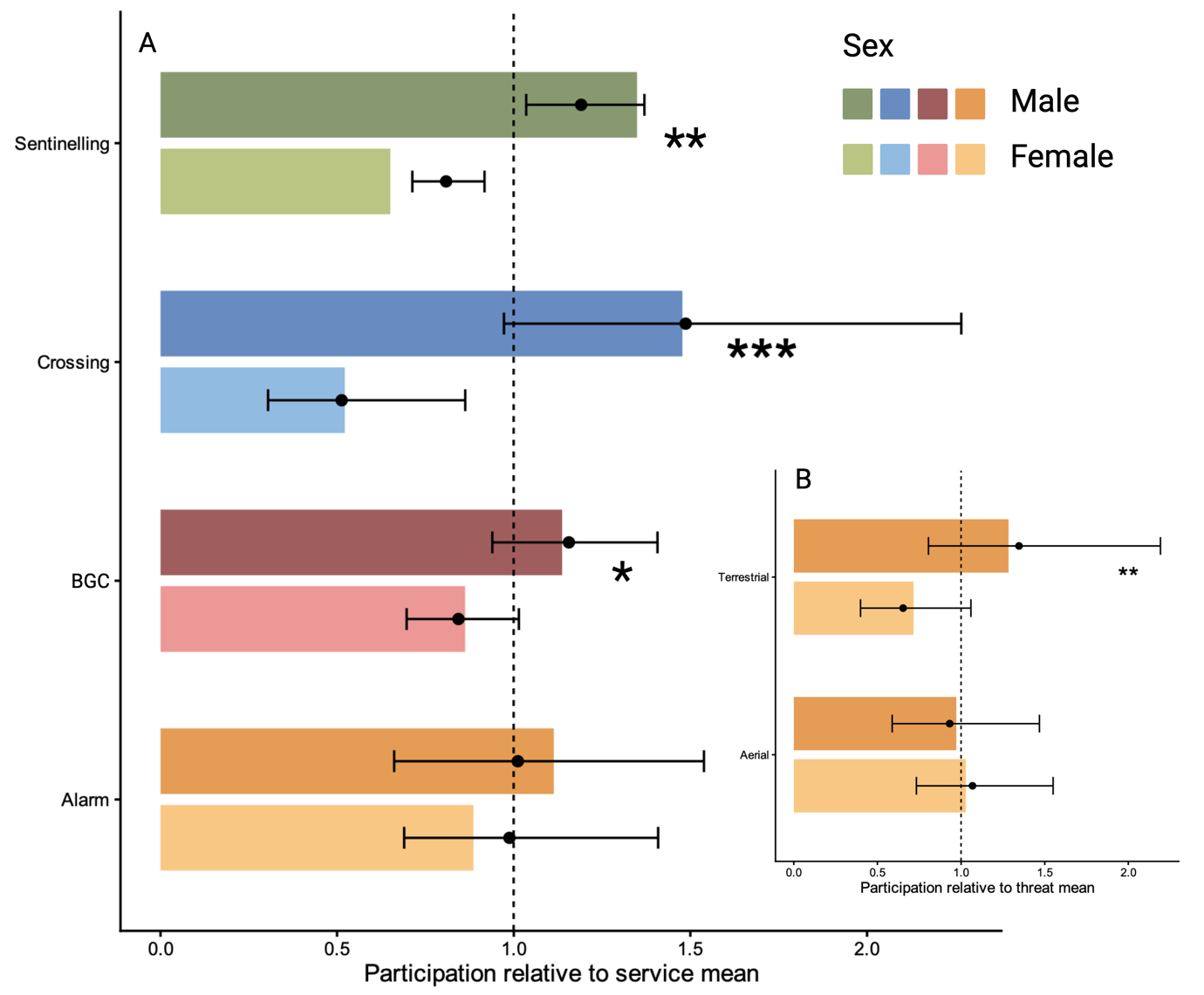

**Fig S1. Participation in cooperative behaviours by males and females relative to the mean level of each service.** Bars show observed mean participation, normalised within each service (time spent sentinelling, leading river crossings, participation in BGC and predator alarms), such that values >1 indicate above-average contribution and values <1 below-average contribution. Points and horizontal error bars represent model predictions (±95% CI). The dashed vertical line at 1 indicates the overall service-specific mean. Males (dark colours) contributed more than females (light colours) across all services, except for alarm calling (A). Notably, male alarm calling was higher than that of females specifically in response to terrestrial predators (B).

Table S2: Statistical models

|  |  | Dates data collection | Model | Null model | | Anova |
| --- | --- | --- | --- | --- | --- | --- |
| Sex differences models | Alarm | 2022-05-10, 2024-05-30 | glmmTMB(participation_alarm ~ Sex * (Threat + ASR + Season) + Group + (1\|AnimalCode) + (1\|EventID), family = binomial, data = model_data_firstmodel_alarm) | *glmmTMB*(participation_alarm ~ Threat + ASR + Season + Group + (1\|AnimalCode) + (1\|EventID), family = binomial, data = model_data_firstmodel_alarm) | | χ²(6) = 10.79, p = 0.095, ΔAIC = 1.2 |
|  | BGC | 2021-10-01, 2025-07-24 | *glmmTMB*(participation_binomial ~ Sex * (BGC_intensity + ASR + Season) + Group + (1 \| AnimalCode) + (1 \| eventID), family = *binomial*(), data = model_data_firstmodel_BGC) | *glmmTMB*(participation_binomial ~ BGC_intensity + ASR + Season + Group + (1 \| AnimalCode) + (1 \| eventID), family = binomial, data = model_data_firstmodel_BGC) | | χ²(7) = 41.01, p < 0.001, ΔAIC = 30 |
|  | Crossing | 2022-04-27, 2025-07-31 | *glmmTMB*(FirstCrosser ~ Sex * (ASR + Season) + Group + (1 \| AnimalCode) + (1\|EventID), family = binomial, data = model_data_model1_crs, offset = *log*(n_males)) | *glmmTMB*(FirstCrosser ~ ASR + Season + Group + (1 \| AnimalCode) + (1\|EventID), family = binomial, data = model_data_model1_crs, offset = *log*(n_males)) | | χ²(5) = 17.61, p = 0.003, ΔAIC =7.6 |
|  | Sentinelling | 2022-05-05, 2025-08-01 | *glmmTMB*(Sentinelling ~ Sex * (ASR + Season) + (1 \| AnimalCode) + Group, ziformula = ~Sex + Season + Group, dispformula = ~ Season + Sex, offset = log(TotalFocalTime), family = nbinom2(), data = model_data_model1_vig) | glmmTMB(Sentinelling ~ ASR + Season + (1 \| AnimalCode) + Group, ziformula = ~ Season + Group, dispformula = ~ Season, offset = log(TotalFocalTime), family = nbinom2(), data = model_data_model1_vig) | | χ²(7)=76.45, p < 0.001, ΔAIC = 63 |
| Male performance models | Alarm | 2022-06-02, 2024-05-30 | glmmTMB(participation_alarm ~ Rank * (Father * Threat + FutureMounts + PastMounts) + Season + zCSI + ASR + Season:FutureMounts + Unhabituated + Group + (1 \| AnimalCode) + (0 + Rank \| AnimalCode) + (1 \| EventID), family = binomial, data = model_data) | *glmmTMB*( participation_alarm ~ Threat + Season + zCSI + ASR + Unhabituated + Group + (1 \| AnimalCode) +(1\|EventID), family = binomial, data = model_data) | | χ²(16) = 28,07, p = 0.03, ΔAIC = 3.39 |
|  | BGC | 2021-10-09, 2024-04-24 | *glmmTMB*( participation_binomial ~ Rank * (Father * BGC_intensity + FutureMounts + PastMounts)+ Season + zCSI + ASR + Season:FutureMounts + Unhabituated + Group + (1 + FutureMounts + Rank + PastMounts\|\| AnimalCode) +(1\|eventID), family = *binomial*(), data = model_data) | *glmmTMB*( participation_binomial ~ BGC_intensity + Season + zCSI + ASR + + Unhabituated + Group + (1 \| AnimalCode) +(1\|eventID), family = binomial, data = model_data) | | χ²(19) = 45.95, p < 0.001, ΔAIC = 8 |
|  | Crossing | 2022-04-28, 2024-07-13 | *glmmTMB*( FirstCrosser ~ Rank * (Father + FutureMounts + PastMounts) + Season + zCSI + ASR+ + Season:FutureMounts + Unhabituated + Group + (1 + FutureMounts + Rank + PastMounts\|\| AnimalCode) +(1\|EventID), family = binomial, data = model_data, offset = *log*(n_males)) | *glmmTMB*( FirstCrosser ~ Season + zCSI + ASR+ + Season + Unhabituated + Group + (1 \| AnimalCode) + (1\|EventID), family = binomial, data = model_data, offset = *log*(n_males)) | | χ²(13) = 19.54, p = 0.11, ΔAIC = 6.46 |
|  | Sentinelling | 2022-05-05 , 2024-06-01 | glmmTMB(Sentinelling ~ Rank * (Father + FutureMounts + PastMounts) + Season + zCSI + ASR + Season:FutureMounts + Unhabituated + Group + (1 + FutureMounts + Rank + PastMounts\|\| AnimalCode), ziformula = ~ Rank + FutureMounts + Season, dispformula = ~ Season + ASR, offset = log(TotalFocalTime), family = nbinom2(), data = model_data) | glmmTMB(Sentinelling ~ Season + zCSI + ASR + Season:FutureMounts + Unhabituated + Group + (1 \|\|AnimalCode), ziformula = ~ Season, dispformula = ~ Season + ASR, offset = log(TotalFocalTime), family = nbinom2(), data = model_data) | | χ²(11) = 35.24, p <0.0001, ΔAIC = 13.3 |
| Mating season model | | *From 04 -01 to 06-30 of years 2023 to 2025.* | Global model | | Best model | |
|  |  |  | *lmer*(*log*((1+number_matings)/days_present) ~ N_AlarmService + VigProp + N_BGCService + N_CrsService # services + Rank + zCSI + + TenureYears # extra variables + year + Unhabituated + (1 \| AnimalCode), # variables to control for data = model_df) | | *lmer*(*log*((1 + number_matings)/days_present) ~ N_BgeService + zCSI + (1 \| AnimalCode), data = model_df) | |

**Table S2**. Overview of the statistical models used to analyse sex differences (models 1), male variance in service provision (models 2), and mating success during the mating season (model 3). For models 1 and 2 each response variable, the table reports the study period, the full (hypothesis-driven) model, the corresponding null model, and likelihood ratio test results comparing full and null models. Test statistics are reported as χ² values with associated degrees of freedom and p-values. For model 3, we report the global model and the best-supported model based on AICc model selection (MuMIn; AICc = 152)

Table S3: Model 1 results

|  | Alarm | | | | BGC | | | | Sentinelling | | | | Crossing | | | |
| --- | --- | --- | --- | --- | --- | --- | --- | --- | --- | --- | --- | --- | --- | --- | --- | --- |
|  | Chisq | Df | p-value | \|βstd\| | Chisq | Df | p-value | \|βstd\| | Chisq | Df | p-value | \|βstd\| | Chisq | Df | p-value | \|βstd\| |
| ASR | 0.283 | 1 | 0.595 | 0.033 | 0.283 | 1 | 0.595 | 0.033 | **9.113** | **1** | **0.003** | **0.082** | 0.002 | 1 | 0.964 | 0.115 |
| BGC_intensity | **182.749** | **2** | **< 0.001** | **0.677** | **182.749** | **2** | **< 0.001** | **0.677** | - | - | - | - | - | - | - | - |
| Group | **22.021** | **3** | **< 0.001** | **0.472** | **22.021** | **3** | **< 0.001** | **0.472** | **8.590** | **3** | **0.035** | **0.203** | **59.061** | **3** | **< 0.001** | **1.687** |
| Season | 3.478 | 3 | 0.324 | 0.162 | 3.478 | 3 | 0.324 | 0.162 | **23.344** | **3** | **< 0.001** | **0.213** | 3.776 | 3 | 0.287 | 0.169 |
| Sex | **5.069** | **1** | **0.024** | **0.027** | **5.069** | **1** | **0.024** | **0.027** | **17.142** | **1** | **< 0.001** | **0.093** | **12.719** | **1** | **< 0.001** | **0.479** |
| Sex:ASR | **8.257** | **1** | **0.004** | **0.104** | **8.257** | **1** | **0.004** | **0.104** | 1.767 | 1 | 0.184 | 0.076 | 0.814 | 1 | 0.367 | 0.120 |
| Sex:BGC_intensity | *4.985* | *2* | *0.083* | *0.048* | *4.985* | *2* | *0.083* | *0.048* | - | - | - | - | - | - | - | - |
| Sex:Season | **23.560** | **3** | **< 0.001** | **0.158** | **23.560** | **3** | **< 0.001** | **0.158** | **16.210** | **3** | **0.001** | **0.166** | 2.954 | 3 | 0.399 | 0.238 |

**Table S3:** Results of Models 1 testing sex differences in cooperative behaviours. For each response variable (alarm calling, between-group conflict participation, sentinelling, and river crossing), the table reports likelihood ratio test statistics (χ²), degrees of freedom, p-values, and absolute standardized effect sizes (|βstd|) for all fixed effects and interactions. Dashes indicate predictors not included in a given model. Statistically significant effects are shown in bold (p < 0.05), and marginal effects are shown in italics (0.05 ≤ p < 0.10).

Table S4: Model 2 results

|  | Alarm | | | | BGC | | | | Sentinelling | | | | Crossing | | | |
| --- | --- | --- | --- | --- | --- | --- | --- | --- | --- | --- | --- | --- | --- | --- | --- | --- |
|  | Chisq | Df | p-value | \|βstd\| | Chisq | Df | p-value | \|βstd\| | Chisq | Df | p-value | \|βstd\| | Chisq | Df | p-value | \|βstd\| |
| Father | 0.099 | 1 | 0.753 | 0.232 | 0.133 | 1 | 0.715 | 0.009 | 0.167 | 1 | 0.683 | 0.081 | 0.003 | 1 | 0.958 | 0.046 |
| Rank | 2.212 | 1 | 0.137 | 1.207 | 0.532 | 1 | 0.466 | 0.022 | 0.426 | 1 | 0.514 | 0.281 | 0.166 | 1 | 0.683 | 0.801 |
| FutureMounts | **9.019** | **1** | **0.003** | **0.851** | **10.878** | **1** | **< 0.001** | **0.148** | **4.662** | **1** | **0.031** | **0.171** | **4.358** | **1** | **0.037** | **0.553** |
| PastMounts | *3.686* | *1* | *0.055* | *0.391* | **4.787** | **1** | **0.029** | **0.184** | 0.720 | 1 | 0.396 | 0.080 | 0.000 | 1 | 0.985 | 0.033 |
| Rank:Father | **5.181** | **1** | **0.023** | **1.112** | 0.919 | 1 | 0.338 | 0.000 | **4.296** | **1** | **0.038** | **0.333** | *3.820* | *1* | *0.051* | *0.674* |
| Rank:FutureMounts | 0.047 | 1 | 0.828 | 0.038 | 0.553 | 1 | 0.457 | 0.045 | **5.017** | **1** | **0.025** | **0.134** | 1.082 | 1 | 0.298 | 0.227 |
| Rank:PastMounts | 0.320 | 1 | 0.572 | 0.092 | 0.038 | 1 | 0.846 | 0.012 | 1.611 | 1 | 0.204 | 0.078 | *3.648* | *1* | *0.056* | *0.397* |
| PredatorThreat | 0.355 | 1 | 0.551 | 0.522 | - | - | - | - | - | - | - | - | - | - | - | - |
| Father:PredatorThreat | 2.410 | 1 | 0.121 | 0.546 | - | - | - | - | - | - | - | - | - | - | - | - |
| Rank:PredatorThreat | 2.285 | 1 | 0.131 | 0.051 | - | - | - | - | - | - | - | - | - | - | - | - |
| Rank:Father:PredatorThreat | 0.742 | 1 | 0.389 | 0.333 | - | - | - | - | - | - | - | - | - | - | - | - |
| BGC_intensity | - | - | - | - | **111.379** | **2** | **< 0.001** | **0.541** | - | - | - | - | - | - | - | - |
| Father:BGC_intensity | - | - | - | - | 0.995 | 2 | 0.608 | 0.143 | - | - | - | - | - | - | - | - |
| Rank:BGC_intensity | - | - | - | - | **6.752** | **2** | **0.034** | **0.408** | - | - | - | - | - | - | - | - |
| Rank:Father:BGC_intensity | - | - | - | - | 2.890 | 2 | 0.236 | - | - | - | - | - | - | - | - | - |
| ASR | 0.071 | 1 | 0.791 | 0.048 | 0.121 | 1 | 0.728 | 0.022 | 2.626 | 1 | 0.105 | 0.112 | **5.370** | **1** | **0.020** | **0.418** |
| zCSI | 1.113 | 1 | 0.291 | 0.205 | 1.504 | 1 | 0.220 | 0.072 | 1.504 | 1 | 0.220 | 0.082 | 0.934 | 1 | 0.334 | 0.162 |
| Season | 1.019 | 3 | 0.797 | 0.200 | *6.264* | *3* | *0.099* | *0.130* | **11.979** | **3** | **0.007** | **0.238** | 0.947 | 3 | 0.814 | 0.171 |
| Group | 4.920 | 3 | 0.178 | 0.654 | **28.911** | **3** | **< 0.001** | **0.626** | 0.932 | 3 | 0.818 | 0.043 | **21.783** | **2** | **< 0.001** | **0.697** |
| Unhabituated | 1.323 | 1 | 0.250 | 0.306 | **11.678** | **1** | **< 0.001** | **0.243** | 0.941 | 1 | 0.332 | 0.065 | 0.475 | 1 | 0.491 | 0.155 |
| FutureMounts:Season | 2.363 | 3 | 0.501 | 0.346 | 0.553 | 3 | 0.907 | 0.044 | **10.536** | **3** | **0.015** | **0.139** | 0.924 | 3 | 0.820 | 0.182 |

**Table S4.** Results of Models 2 testing predictors of male investment in cooperative behaviours. For each response variable (alarm calling, between-group conflict participation, sentinelling, and river crossing), the table reports likelihood ratio test statistics (χ²), degrees of freedom, p-values, and absolute standardized effect sizes (|βstd|) for all fixed effects and interactions. Dashes indicate predictors not included in a given model. Statistically significant effects are shown in bold (p < 0.05), and marginal effects are shown in italics (0.05 ≤ p < 0.10).

Table S5: Individual differences models

| Service | Anova | ΔAIC | Among-male variance (SD) | Among-event variance (SD) |
| --- | --- | --- | --- | --- |
| Alarm | χ²(1) = 9.2, p = 0.002 | 7.21 | 0.66 (0.81) | <0.001(<0.01) |
| BGC | χ²(1) = 62.78, p <0.0001 | 60.8 | 0.23 (0.48) | 0.29 (0.53) |
| Crossing | χ²(1) = 5.05, p =0.024 | 5.05 | 0.54 (0.73) | <0.001(<0.0001) |
| Sentinelling | χ²(1) = 5.82, p =0.016 | 5.09 | 0.065 (0.25) | - |

**Table S5**. Evidence for consistent individual differences in male service provision. For each service, we compared the final research model (Table S3), excluding random slopes, with an otherwise identical model that omitted male identity (AnimalCode) as a random effect. All models retained the same fixed effects and event-level random effects where applicable. ΔAIC and likelihood-ratio tests (χ²) quantify the improvement in model fit associated with including male identity. Male identity variance represents the estimated among-male variation in service provision after accounting for other predictors in the model, with the standard deviation in parenthesis, same for event ID when applicable. Larger values indicate greater variation attributable to male identity or event-level differences, respectively. In all services, male identity explained significant variation, indicating persistent differences among males in their propensity to provide a given service.

|  | Group | Number of males (median ± SD) | | Group size (median ± SD) | | Number of focals | | Number of predator events | | Number of between-group conflict events | | | Number of Crossing events |
| --- | --- | --- | --- | --- | --- | --- | --- | --- | --- | --- | --- | --- | --- |
|  |  |  |  |  |  | Males | Females | Aerial | Terrestrial | Minor | Medium | High |  |
| Sex differences | AK | 2 | 0.76 | 23 | 3.01 | 286 | 542 | 19 | 8 | 45 | 103 | 80 | 3 |
|  | BD | 10 | 3.47 | 70 | 13.08 | 849 | 457 | 67 | 10 | 54 | 116 | 61 | 78 |
|  | KB | 4 | 1.07 | 22 | 1.00 | 249 | 460 | 20 | 12 | 26 | 41 | 16 | 31 |
|  | NH | 8 | 1.32 | 45 | 8.18 | 575 | 886 | 58 | 18 | 65 | 99 | 62 | 71 |
| Male performance | AK | 3 | 0.47 | 23 | 1.64 | 192 | 0 | 5 | 6 | 36 | 77 | 54 | NA |
|  | BD | 10 | 3.37 | 72 | 10.85 | 642 | 0 | 28 | 11 | 49 | 95 | 44 | 44 |
|  | KB | 3 | 0.48 | 22 | 1.07 | 141 | 0 | 7 | 3 | 22 | 31 | 12 | 15 |
|  | NH | 8 | 0.82 | 47 | 5.3 | 348 | 0 | 13 | 7 | 59 | 80 | 51 | 16 |
| Mating season success | year | 2023 | | | | 2024 | | | | 2025 | | | |
|  | Group | Number of males | | Group size | | Number of males | | Group size | | Number of males | | Group size | |
|  | BD | 17 | | 72 | | 11 | | 43 | | 5 | | 24 | |
|  | KB | 3 | | 22 | | 4 | | 23 | | 2 | | 18 | |
|  | NH | 10 | | 47 | | 7 | | 29 | | 3 | | 22 | |

Table S6: Group composition

**Table S6**. Summary of the analysed data and group composition across study groups and analyses. Differences in sampling periods across behaviours reflect constraints imposed by changes in data collection protocols; in all cases, the maximum amount of data was included without altering protocols.
